# Slow neural oscillations explain temporal fluctuations in distractibility

**DOI:** 10.1101/2022.10.04.510769

**Authors:** Troby Ka-Yan Lui, Jonas Obleser, Malte Wöstmann

## Abstract

Human environments comprise various sources of distraction, which often occur unexpectedly in time. The proneness to distraction (i.e., distractibility) is posited to be independent of attentional sampling of targets, but its temporal dynamics and neurobiological basis are largely unknown. Brain oscillations in the theta band (3 – 8 Hz) have been associated with fluctuating neural excitability, which is hypothesised here to explain rhythmic modulation of distractibility. In a pitch discrimination task (N = 30) with unexpected auditory distractors, we show that distractor-evoked neural responses in the electroencephalogram and perceptual susceptibility to distraction were co-modulated and cycled approximately 3 – 5 times per second. Pre-distractor neural phase in left inferior frontal and insular cortex regions explained fluctuating distractibility. Thus, human distractibility is not constant but fluctuates on a subsecond timescale. Furthermore, slow neural oscillations subserve the behavioural consequences of a hitherto largely unexplained but ever-increasing phenomenon in modern environments – distraction by unexpected sound.

## Introduction

Selective attention enables humans to focus on relevant information at the expense of distraction. The brain prioritizes representations of relevant events while filtering out task-irrelevant distractors (Desimone & Duncan, 1995; Picton et al., 1971). Recent research posited that distractor processing is not merely collateral to attentional sampling of targets but may follow its own dynamics (Schneider et al., 2018; Wöstmann et al., 2019, 2020). The behavioural detriments induced by different kinds of distractors (i.e., *distraction*) and the neuro-cognitive mechanisms that counteract distraction (i.e., *suppression*) have been studied in some detail (Bonnefond & Jensen, 2012; Geng & DiQuattro, 2010; van Moorselaar et al., 2020; Weisz et al., 2020; Wöstmann et al., 2019). However, the temporal dynamics and the neurobiological basis of the proneness to distraction (i.e., *distractibility*) are largely unknown.

Distractibility has long been neglected in the theoretical formulation of rhythmic attention. Originally assumed to be static (Posner et al., 1980), the attentional spotlight was proposed to be blinking at a subsecond time scale in a theta-like rhythm (i.e., 3 – 8 Hz) (Buschman & Kastner, 2015; Fiebelkorn & Kastner, 2019). Behaviourally, it is manifested via the waxing and waning of behavioural performance in target selection (Fiebelkorn et al., 2013; Ho et al., 2017; Kubetschek & Kayser, 2021; Landau & Fries, 2012) or working memory (Schmid et al., 2022; ter Wal et al., 2021) performance at similar frequencies. However, the temporal dynamics outside of the attentional spotlight are not well understood. While previous research studied how distractibility unfolds on relatively long temporal scales of minutes (i.e., during an experimental session (Forster & Lavie, 2014)) or years (i.e., across stages of development (Campbell et al., 2012; Kannass et al., 2006)), we found preliminary evidence for fluctuating distractibility on shorter timescales following rhythmic presentation of auditory targets (Wöstmann et al., 2020). To isolate distractibility dynamics from rhythmic entrainment or preparatory suppression, we here employ a design that uses non-rhythmic stimuli and distractors that occur unexpectedly.

A central prediction of rhythmic attention is that the phase of slow neural oscillations explains fluctuations in behaviour (VanRullen, 2016). The prediction is based on the notion that rhythmic attention arises from the periodic excitability of the attention-related brain network (Fiebelkorn & Kastner, 2019; VanRullen, 2016). In the human brain, theta neural phase (3 – 8 Hz) is assumed to reflect moment-to-moment changes in neural excitability (Lakatos et al., 2005). Theta phase in brain regions beyond sensory cortices, such as fronto-parietal regions and the hippocampus, has been associated with fluctuations in target detection (Helfrich et al., 2018) and working memory encoding (Rutishauser et al., 2010; Siegel et al., 2009), respectively. While previous research has related distractibility to supra-modal regions in frontal (Chao & Knight, 1995; Wais et al., 2012) or parietal (Kanai et al., 2011) cortex, it is unclear whether and in which networks the momentary neural dynamics may subserve the waxing and waning of distractibility.

Here, we ask if the brain spontaneously alternates between states of higher and lower distractibility and whether such fluctuations have the potency to explain behavioural consequences of distraction. If so, we would expect to observe a brain-behaviour relation between the pre-distractor brain state and the distractor-induced detriment in task performance. To this end, we employed a pitch discrimination task wherein an auditory distractor could occur at variable and unexpected times in-between two target tones. We probed this research question in the auditory modality as temporal information is especially important to auditory attentional selection (Shamma et al., 2011). During the task, participants had to identify whether the two target tones were the same or different in pitch (Fig. 1A). The distractor was a fast-varying, 25-Hz modulated sequence of tones that differed in pitch, which allowed us to extract its evoked 25-Hz neural response (Ding & Simon, 2009).

**Fig 1.**
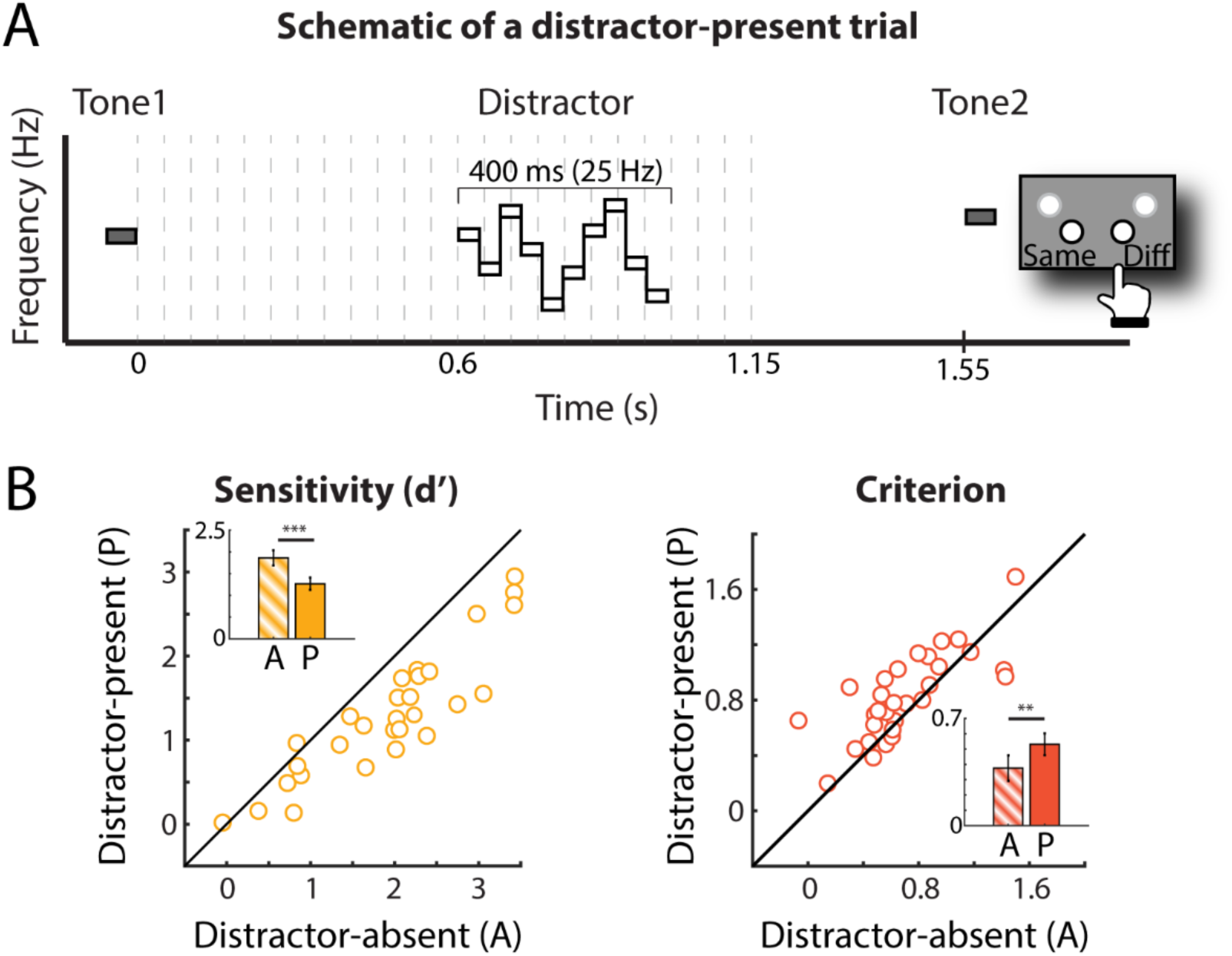
Experimental design and behavioural results. A) Schematic of a distractor-present trial. Participants were instructed to indicate whether the two target tones (grey) were the same (probability = 50%) or different (higher, probability = 25%; or lower, probability = 25%) in pitch. A 10-tone-pip distractor sequence (white) with a 25-Hz temporal structure (i.e., 40-ms tone-pip duration; total duration 400 ms) was presented at one of the 24 distractor onset times (dashed lines). In distractor-absent trials, no distractor was presented. B) Behavioural results comparing distractor-present and -absent conditions. Coloured circles indicate single-subject data. Insets show bar graphs of perceptual sensitivity (left panel) and criterion (right panel) for distractor-present (solid bar) and distractor-absent (gradient bar) conditions, respectively. Error bars show ±1 SEM. ** p < .01. *** p < .001.

We used behavioural sensitivity and distractor-evoked neural response as the behavioural and neural proxies of distraction. Behaviourally, perceptual sensitivity was calculated as an indirect measure of distraction: The more distracted, the lower the sensitivity in pitch discrimination should be. Neurally, we extracted the amplitude of the distractor-evoked event-related potential (ERP) at 25 Hz, which corresponded to the modulation rate of the frequency-modulated distractor tone sequence. Although these post-distractor measures may not solely reflect distractibility but also other distractor-related processes (e.g., suppression), we should observe temporal fluctuations of the two measures if distractibility exhibits temporal dynamics. If there is a brain-behaviour relation in the temporal fluctuations in distraction, we should also observe temporal co-fluctuations between the two measures. Furthermore, we aimed to unveil the neural origins of the distractibility dynamics. If a brain region is involved in the momentary changes in distractibility, we should observe a relationship between the pre-distractor phase of neural oscillation in that region and the fluctuations in behavioural performance.

A total of 17,280 behavioural and neural responses in the electroencephalogram (EEG) in N=30 participants revealed that behavioural sensitivity and distractor-evoked neural responses fluctuated in sync across distractor onset times in ∼3 – 5 cycles per second. Critically, pre-distractor theta phase in left inferior frontal and insular cortex regions explained behavioural performance fluctuations. These effects were absent in trials without distractors, reinforcing their specificity to distractor-related neural processing.

## Material and Methods

### Participants

Thirty participants (20 females, 10 males; mean age = 23.67, SD = 3.56) took part in the EEG experiment. They provided written informed consent and were compensated by either €10/hour or course credit. Participants were right-handed according to the Edinburgh Handedness Inventory (Oldfield, 1971) (mean score = 92), with self-reported normal hearing, normal or corrected-to-normal vision, and no psychological or neurological disorders. All procedures of the current study were approved by the ethics committee of the University of Lübeck.

### Stimuli and Procedure

Participants performed a pitch discrimination task wherein they decided whether the first (tone 1) and the second (tone 2) target tones in a trial were the same or different in pitch. Prior to the experiment, they were instructed to answer as accurately and as fast as possible. The target tones were 75 ms long pure tones with 5 ms rise and fall periods. In each trial, the frequencies of tone 1 were randomly selected between musical note A#3 (233 Hz) and G#5 (830.6 Hz), while that of tone 2 was either the same (50%) or different (higher or lower, 25% each) in frequency compared to tone 1.

The pitch difference between tone 1 and tone 2 was titrated for each participant with an adaptive task (see below). The offset-to-onset interval between tone 1 and tone 2 was 1550 ms. Each distractor stimulus comprised 10 consecutive pure tones with 40 ms duration (400 ms in total). The frequencies of the pure tones in each distractor stimulus were randomly selected among the 12 tones between A#3 and G#5 with whole tone steps (A#3, C4, D4, E4, F#4, G#4, A#4, C5, D5, E5, F#5, and G#5), with the constraint that there would be no repetition between consecutive tones. Each of the 12 tone frequencies appeared at each of the 10 positions with equal probability across trials.

In-between the two target tones, a distractor was presented in 50% of trials (distractor-present condition) and no distractor was presented in the remaining trials (distractor-absent condition). The inclusion of distractor-absent trials serves two purposes. First, we could verify that the distractors had the potency to distract by comparing behavioural performance for distractor-present versus distractor-absent trials (Wöstmann et al., 2022). Second, participants could not anticipate whether or when a distractor would occur in a given trial, which eliminated potential effects of such anticipation on behavioural performance (Grabenhorst et al., 2021) or pre-stimulus neural activity (Dürschmid et al., 2018; Herbst et al., 2022; Stefanics et al., 2010). The distractor was a tone sequence which consisted of 10 40-ms tone pips, thus creating a 25-Hz temporal structure (total duration: 400 ms).

In the distractor-present condition, the distractor was presented at one of 24 distractor onset times (0 ms to 1150 ms, 50-ms steps, relative to the offset of tone 1), which was selected at random on each trial. The distractor-absent trials were randomly assigned to the 24 distractor onset times. Specifically, for each distractor-absent trial, a distractor onset time was assigned as in the distractor-present trial. A distractor is however not presented during stimulus presentation in the distractor-absent condition. Time 0 in a distractor-absent trial, therefore, refers to the time when a distractor would have been presented in that trial.

After the offset of target tone 2, participants had a 2000 ms response time window. After the presentation of tone 2, a prompt is shown on the screen asking if the two target tones were the same or different in pitch (“same” or “different”?). Participants were only allowed to respond after the presentation of tone 2. Any button press beforehand was thus not recorded. To avoid potential temporal predictability effects of the onset of the next trial, the inter-trial intervals were randomly selected from a truncated exponential distribution (mean = 1460 ms), ranging between 730 and 3270 ms.

The trial order was pseudo-randomized with no repetition in probe tone frequency and distractor onset for any two consecutive trials. In total, there were 12 trials for each unique condition (distractor-present/absent x distractor onset x same/different target pitch) and 1152 trials for the whole experiment. All auditory materials were presented via Sennheiser headphones (HD 25-1 II). Responses were made using a response box (The Black Box Toolkit). The assignment of buttons to the response options (“same” or “different”) was counterbalanced across participants. Stimuli were presented via Matlab (MathWorks, Inc., Natick, USA) and Psychtoolbox(Brainard, 1997). The auditory stimuli were presented at approximately 70 dB SPL.

### Adaptive Staircase Procedure

Prior to the main experiment, each participant’s threshold for the pitch discrimination task was titrated using an adaptive staircase procedure, implemented in the Palamedes toolbox (Prins & Kingdom, 2018) for Matlab. For the initial 11 participants, the threshold was titrated to an approximate accuracy of 70.7%. As the overall accuracy was relatively high even after the adaptive staircase procedure for these 11 participants (mean = 79.59%, SD = 10.43%), the final 16 participants performed an adaptive procedure altered to yield approximately 65% accuracy instead. Due to technical issues, performance of the remaining three participants was tracked at 35% accuracy. As all relevant statistical analyses in the present study are within-subject, and as paired t-tests (2-tailed) comparing the behavioural performance between distractor-absent and distractor-present conditions were significant with (*t_29_* = 8.11, *p* < .001) and without (*t_26_* = 9.41, *p* < .001) these participants, their data were included in the final analysis.

Each participant went through the adaptive staircase procedure two to three times, depending on the stability of the tracked threshold. The range of frequencies used in the adaptive task was the same as that used in the main experiment (i.e., 233 Hz to 830.6 Hz). There were in total 30 trials for each run of the adaptive staircase procedure with an initial pitch difference of 100 cents (i.e. 1 semitone) between tone 1 and 2. The minimum and maximum pitch difference possible in the task was 2 cents and 2000 cents, respectively. For the procedure which tracked performance at ∼70.7%, a two-down one-up procedure was used. Specifically, the pitch differences would decrease in steps of 10 cents if participant responded correctly (i.e., different), or increase in steps of 10 cents if participant responded incorrectly (i.e., same) for 2 consecutive trials. For the procedure which tracked performance at ∼65% procedure, the pitch differences would decrease in steps of 7 cents if participant answered correctly or increase in steps of 13 cents if they answered incorrectly. The pitch difference used in the main experiment was calculated by averaging the final 10 trials in the tracking run which converged to the most stable threshold, determined by visual inspection, in the ∼70.7% procedure. The same procedure was used to average the final 6 trials in the ∼65% procedure.

Overall accuracy averaged across all participants in the actual experiment was 73.58% (SD = 12.12%). For participants tracked to 65 %, the final threshold ranged from 22.5 to 269.2 cents. For participants tracked to 70.7%, the final threshold ranged from 35 to 180 cents. The average threshold across participants was 95.4 cents and the standard deviation 77.4 cents. Participants with a higher level of tracked accuracy performed better in the main experiment (*t_28_* = 3.66, *p* = .001).

### Behavioural Data Analysis

To understand how distractors affect pitch discrimination performance in the framework of signal detection theory, we calculated sensitivity (d’) and criterion (c) separately for distractor-present and -absent conditions, using the Palamedes toolbox (Prins & Kingdom, 2018) and the following formulas:

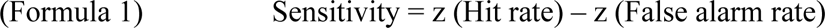

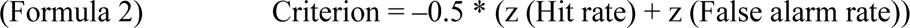

Hit rate was defined as the “different” response when the two tones were different in pitch, and false alarm rate the “different” response when the two tones were the same in pitch. Extreme values (0 or 1) of Hit rate or False alarm rate were adjusted (Macmillan & Kaplan, 1985): A rate of 0 was adjusted by dividing 1 by the number of trials multiplied by 2; while a value of 1 was adjusted by subtracting the same value from 1. Paired samples t-tests (2-tailed) were used to compare sensitivity and criterion in distractor-present versus-absent conditions.

To study the modulation of distractor onset times on behavioural measures in the distractor-present condition, sensitivity for each distractor onset time was calculated, resulting in a behavioural time course as a function of distractor onset time for each individual participant (see Fig. S1 & Fig. S2).

### EEG Recording and Pre-processing

The experiment was conducted in an electrically shielded sound-attenuated room. A modified 10-20 international system with 64 Ag/Ag-Cl electrodes was used to record the EEG with a sampling rate of 1000 Hz (actiCHamp, Brain Products, München, Germany). The EEG recordings were band-pass filtered online from direct current (DC) to 280 Hz. TP9 was used as the online reference and FPz as the ground electrode. Impedances were kept below 20 kOhm for all but one participant.

Matlab R2018a (MathWorks, Inc., Natick, USA) and the Fieldtrip toolbox (Oostenveld et al., 2011) were used to pre-process and analyse EEG data. The continuous EEG data were filtered (high-pass, 1 Hz; low-pass, 100 Hz) before they were segmented into epochs (−2 to 2.5s) time-locked to tone 1 onset. Independent component analysis (ICA) was used to identify and reject components corresponding to artefacts such as eye blinks, eye movements, and muscle activity (average percentage of components removed = 26.46%, SD = 8.89%). Afterwards, EEG data were re-referenced to the average of all electrodes. Epochs with amplitude changes >160 microvolts were rejected (average percentage of epochs removed = 1.35%, SD = 2%).

To obtain distractor-evoked neural responses, data were re-epoched to the onset of the distractor (−1 to 1 s) with a 200ms baseline period before distractor onset (i.e., −200 to 0ms). Epochs belonging to the same conditions (distractor-present/absent) and distractor onset time (0 – 1150ms, 50-ms steps) were then averaged into ERP waveforms. The spectral amplitude of distractor-evoked responses at 25 Hz, which corresponds to the temporal structure of the distractor, was extracted using FFT on the ERP waveform in the time window from 0 to 520ms after distractor onset. The frequency spectrum of the distractor-evoked ERP waveform shows a distinct peak of the spectral amplitude at 25 Hz (Fig. S3; Donoghue et al., 2020). Spectral amplitude was averaged across electrodes F1, Fz, F2, FC1, FCz, and FC2. For each participant, the 24 spectral amplitudes, corresponding to the 24 distractor onset times, resulted in a neural time course of distractor processing as a function of distractor onset time (see Fig. S1 & Fig. S2).

We chose the spectral amplitude of distractor-evoked responses at 25 Hz instead of N1 amplitude as the neural measure of distraction for two main reasons. First, the distractor-evoked ERP amplitude at 25 Hz reflects the evoked neural responses to all 10 distractor tones in the distractor tone sequence, which would have a higher SNR compared to the N1 amplitude that essentially reflects the neural response to the distractor-sequence onset. Second, the difference in the N1 component across distractor onset time is not limited to amplitude, but also apply to latency and morphology (see Fig. S4). Modulation of the N1 component may reflect multiple components, such as deviance detection (Wang et al., 2008) or the encoding of tone 1 for trials with early distractors. Hence, distractor-evoked responses at 25 Hz were deemed a more appropriate and more specific neural proxy of distraction in the current study.

Distractor-evoked inter-trial phase coherence (ITPC) was also calculated across frequencies (1 – 10 Hz, 1-Hz steps) and time windows (−0.2 – 0.7 s, 0.05-s steps) for each electrode. First, Fourier coefficients were calculated (using windows with a fixed length of 0.5 s; hanning taper). Then, the complex Fourier coefficients were divided by their magnitude and averaged across trials. ITPC was calculated by taking the absolute value (i.e., magnitude) of the average complex coefficient.

### Modulation of neural and behavioural measures by distractor onset time

To test whether and how distractor onset time modulates neural and behavioural measures, we used linear mixed-effect models with sine- and cosine-transformed distractor onset time, similar to Wöstmann et al. 2020 (Wöstmann et al., 2020). This method outperforms other methods for studying the phasic modulation of behavioural and neural responses (Zoefel et al., 2019) and has also been used previously (Wöstmann et al., 2020) to extract temporal fluctuations in the vulnerability of working memory to distraction. A quadratic trend was observed in the behavioural time course in Fig. 2A as the earliest and latest distractors were most distracting due to their temporal proximity to the target tones. For time courses of sensitivity and spectral amplitude of the distractor-evoked ERP at 25 Hz separately, we first subtracted the individually fitted quadratic trend (computed with the polyfit function in Matlab) from the original time course for each participant (see Fig. S1 & Fig. S2) as the quadratic trend was not of interest in the current study (Huang et al., 2015).

**Fig 2.**
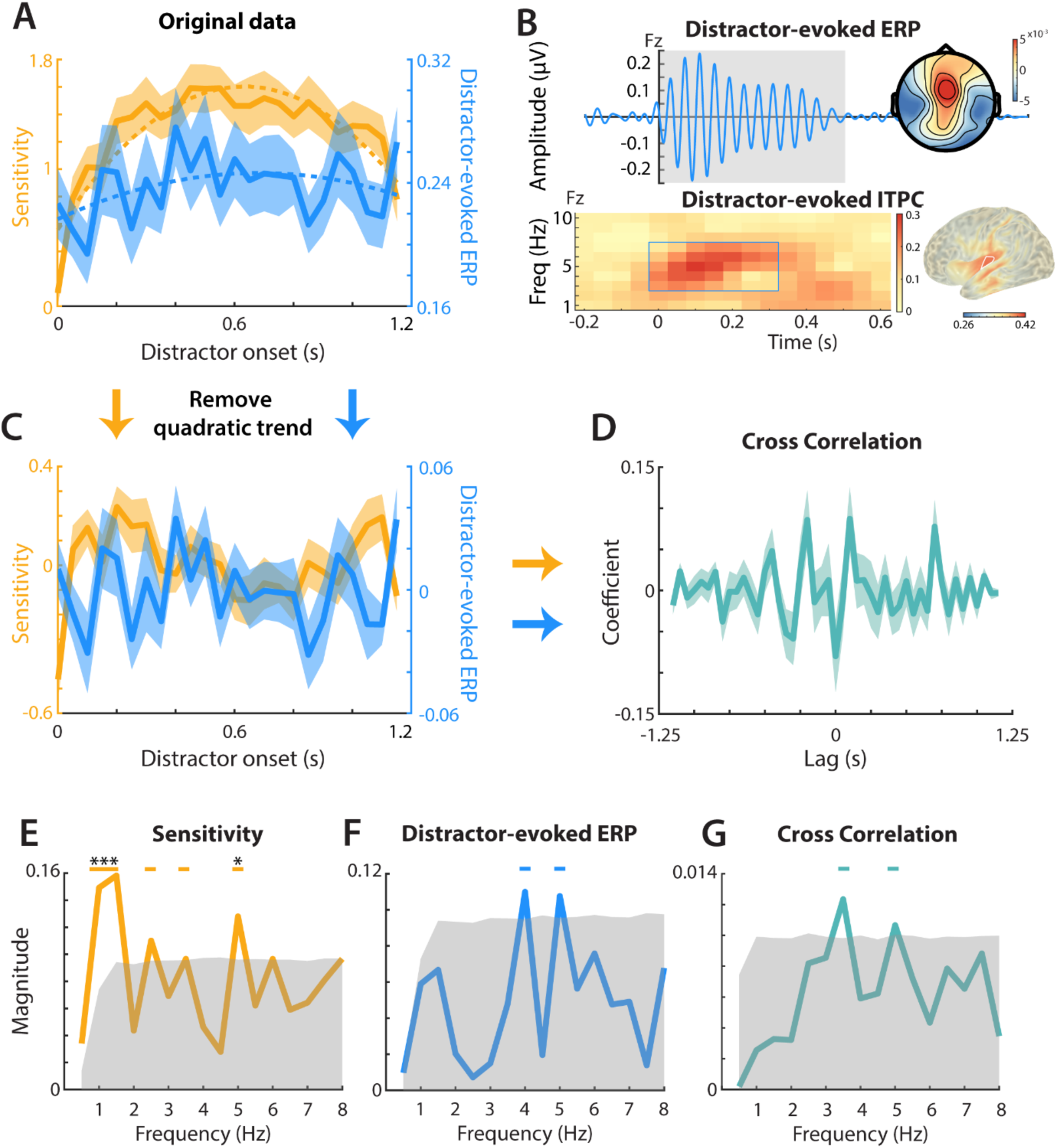
The analyses of behavioural and neural time courses by distractor onset time. A) Average sensitivity (yellow solid line) and 25-Hz amplitude of the distractor-evoked event-related potential (ERP; blue solid line) across distractor onset times. Shaded areas show ±1 SEM across participants. Dashed lines show respective quadratic trends. B) Top panel: Distractor-evoked ERP waveform averaged across all distractor onset times at electrode Fz (20 – 30 Hz bandpass filtered for visualization purpose). Shaded grey area marks the time window used to extract the 25-Hz amplitude of the distractor-evoked ERP. Inset shows the scalp map of the 25-Hz amplitude of the distractor-evoked ERP (derived via an FFT on the distractor-evoked ERP waveform). Bottom panel: Distractor-evoked inter-trial phase coherence (ITPC) from 1 – 10 Hz and from −0.2 s – 0.6 s at Fz. Brain surface shows the ITPC values (frequencies: 3 – 7 Hz; time window: 0 – 0.3 s) in source space, which reflects the auditory response to the distractor. White outline indicates top 1% voxels with largest ITPC values. C) Detrended time courses of behavioural and neural outcome measures. Shaded areas show ±1 SEM across participants. D) Solid line shows average correlation coefficients, derived by averaging single-subject cross-correlations of sensitivity and distractor-evoked ERP time courses, as a function of temporal lags. Shaded area shows ±1 SEM across participants. E-G) Spectral magnitude across frequencies (0.5 – 8 Hz, 0.5-Hz step) for (E) detrended sensitivity, (F) distractor-evoked ERP, and (G) the cross-correlation between the two. Shaded areas show the 95^th^ percentile of the permutation distribution generated from 5,000 permutations. Horizontal lines show statistical significance when comparing the spectral magnitude against the 95^th^ percentile of the permutation distribution before FDR correction. Asterisks show the statistical significance that remained significant after FDR correction.

Then, we designed sine- and cosine-transformed distractor onset time vectors using the following formulas,

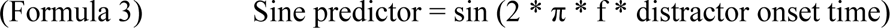

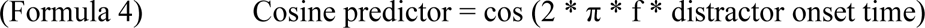

Where f denotes the frequency of interest (0.5 – 8 Hz, 0.5-Hz steps). Next, we regressed the detrended sensitivity and spectral amplitude of ERP time courses on sine and cosine predictors using linear mixed models (using the fitlme function in Matlab) for each frequency of interest using the following formulas:

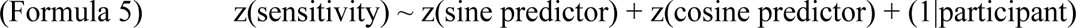

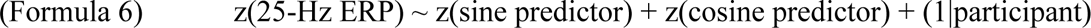

Spectral magnitude for each frequency was computed by taking the square root of the sum of squared beta coefficients of sine and cosine predictors:

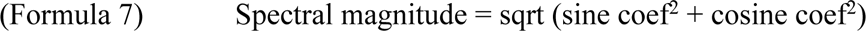

Statistical significance of the spectral magnitude was determined by comparing the spectral magnitude of the empirical data with the 95^th^ percentile of a permutation distribution, which was generated by shuffling the original behavioural/neural time course and performing the same analysis for 5,000 times.

To test whether sensitivity and spectral amplitude of the distractor-evoked ERP at 25 Hz are co-modulated, for each participant, cross-correlation coefficients across time lags of the two signals were obtained. Specifically, the cross-correlation coefficients per participant were obtained from the z-transformed detrended behavioural sensitivity time course and distractor-evoked ERP time course (Two vectors; yellow and blue lines in Fig. 2C). We used the matlab function xcorr(zscore(detrended sensitivity), zscore(detrended distractor-evoked ERP)) to obtain the cross-correlation coefficients. Again, we ran a similar linear mixed model as explained above, but this time with sine- and cosine-transformed time lags as predictors and used the correlation coefficients from the cross-correlation as the outcome measure. Spectral magnitude was obtained using formula 7 and statistical significance with the same permutation method mentioned above.

Alternative to the shuffling-in-time approach, the frequency spectra of the behavioural and neural time courses in the distractor-present condition can be contrasted against those in the distractor-absent condition to test for the temporal fluctuations in the behavioural and neural proxies of distraction. For behavioural sensitivity, distractor-evoked ERP, and cross-correlation time courses, we additionally compared the spectral magnitude between distractor-present and -absent conditions with correction for multiple comparisons (i.e., FDR correction).

### Phasic modulation of behavioural sensitivity

To explore the role of pre-distractor neural dynamics on the pitch discrimination performance, we examined whether pre-distractor oscillatory phase relates to behavioural sensitivity. To this end, we examined the quadratic fit of sensitivity as a function of neural phase in source space.

First, we implemented the source analysis using the Fieldtrip toolbox. First, a standard volume conduction model and standard electrode locations were used to calculate the leadfield matrix with 10-mm resolution. We applied the linearly constrained minimum variance (Van Veen et al., 1997) (LCMV) beamformer approach on the 10 Hz lowpass filtered data centred around distractor onset (−1 to 1s). We calculated a common filter including all trials by calculating the covariance matrix estimates. There were in total 2,015 source locations inside the brain.

Second, a quadratic fit analysis resolved by frequency and time probed the spectral and temporal specificity of the phasic modulation of perceptual sensitivity. To obtain trial-wise phase values for each source location, the following procedure was implemented for each trial in each source location: First, the single-trial EEG time course was projected into the source space using the common filter. Then, a sliding window (0.4s duration; moving in 50-ms steps from –0.3 to +0.3s relative to distractor onset) was employed to transform the data into the frequency domain (using FFT). Note that the time point of the sliding window refers to the mid-point of each time window. For instance, the time window centred at −0.3 included data from −0.5 to −0.1 s. The respective phase value of each frequency (2.5 – 8 Hz in 0.5-Hz steps) was then calculated using the *angle* function in MATLAB. The phase values of all trials were binned into 9 bins of equal size, ranging from −pi to pi, followed by a calculation of sensitivity for each bin. The quadratic fit of sensitivity across phase bins was estimated using the *polyfit* function (order = 2) in MATLAB. As a result, we obtained a quadratic fit index for each source location, frequency, and time of interest.

The choice of the quadratic fit analysis rather than Kullback-Leibler divergence (i.e., KL divergence), which captures potential patterns beyond the quadratic trend, is supported by the following reasons: First, the quadratic fit analysis is more specific than KL divergence as it only captures cyclic modulations of sensitivity by neural phase. KL divergence captures the deviation in the distribution of sensitivity by neural phase regardless of the pattern of the phase dependence (see Fig. S5).

Second, when doing the quadratic fit analysis on one fixed time window only, the quadratic fit may fail to capture the phasic relationship that manifests as a sine wave function. We circumvented this potential caveat by running a time-resolved quadratic fit analysis. By shifting the analysis window in time, we are able to capture cyclic modulations with different phase shifts.

We used a source-level cluster-based permutation test (Maris & Oostenveld, 2007) to find significant clusters in voxel-frequency-time space that would exhibit phasic modulation of sensitivity. Dependent-samples t-tests were used to contrast quadratic fit coefficients against zero, followed by clustering of adjacent bins with significant effects in voxel-frequency-time space. To derive cluster p-values, summed t-values in observed clusters were tested against 5,000 permutations with shuffled condition labels (two-tailed).

To demonstrate that the significant cluster found in the above analysis does not primarily originate from auditory cortex, we localised, for comparison, the distractor-evoked inter-trial phase coherence (ITPC) at 3 – 7 Hz, strongly assumed to emerge at least to large degrees from the supratemporal plane and auditory cortex (Koerner & Zhang, 2015; Mayhew et al., 2010; Oya et al., 2018), with the following procedure for each voxel: For each trial, we projected the time series EEG data into source space using the same common filter as in the analysis on the phasic relationship with behaviour. Then, we transformed the source-projected data (0 – 300 ms after distractor onset) to the frequency domain using FFT. The same calculation as on the sensor level was used to calculate the ITPC for each frequency. ITPC across frequencies 3 – 7 Hz were then averaged to obtain one distractor-evoked ITPC value for each voxel.

One may ask how the dynamics in distractibility relate to rhythmic attentional sampling. As the current study mainly focuses on distractibility dynamics, we did not use the dense sampling approach to vary target onset time. Hence, we cannot test for the endogenous dynamics in attentional sampling. Nevertheless, if the phase dependence of distractibility is indeed an anti-phase version of the phase dependence of attentional sampling, we should observe a quadratic phasic relationship between the phase of theta neural oscillations before tone 2 onset and behavioural sensitivity, which would have an inverse pattern compared to the quadratic trend found for pre-distractor neural phase. Furthermore, the phasic relationship should be observed in the same brain regions and frequencies as the relation between neural phase and distractibility. To test this hypothesis, we ran the same quadratic fit analysis with the neural phase before tone 2 onset for distractor-present and -absent trials, separately. We restricted the analysis to the frequencies and time windows where we found the significant positive cluster in the main analysis (Fig. 3C). We ran a cluster permutation test within the frequencies and time windows with the same parameters as the main analysis on the quadratic fits.

**Fig 3.**
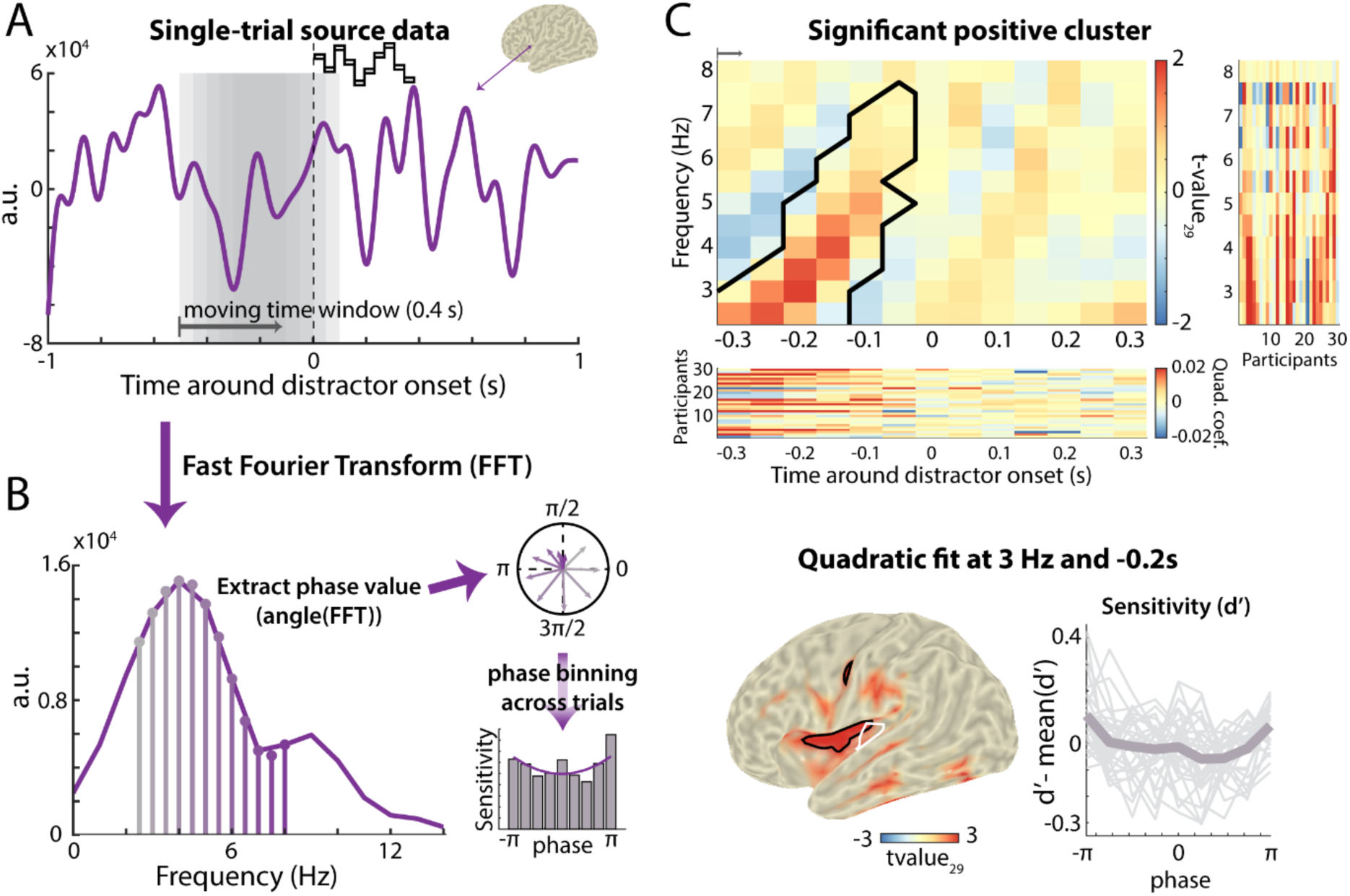
Cluster-based permutation test results on the relationship between neural phase and behavioural fluctuations. A-B) Illustration of the source-level analysis. A) Example of a single-trial source-projected EEG time course. The moving window (grey) was used to transform segments of the data into the frequency domain using FFT. The first grey window corresponds to the first time window used in the time-resolved analysis (i.e., −0.5 to −0.1 s). B) Spectral representation of the data segment in (A). Phase values across frequencies were extracted and trials were binned according to their phase values into 9 phase bins for each frequency, time window, and source location. Bar graph shows exemplary sensitivity values calculated from the trials sorted by phase bin. A quadratic trend was fitted to the sensitivity values across phase bins (purple solid line). C) Results of a cluster-based permutation test, which tested quadratic fits in time-frequency-source space against zero. Top panel shows the t-values (df = 29) across frequencies and time windows, averaged across all the voxels belonging to the significant positive cluster. The black contour indicates the positive significant cluster. Right column shows individual participants’ quadratic coefficients for each frequency, collapsed across the time windows included in the significant cluster. Bottom row shows individual participants’ quadratic coefficients across time windows, collapsed across frequencies and voxels included in the significant cluster. Bottom left panel shows the cluster peak effect (3 Hz; −0.2 s), which resides mainly in left inferior frontal cortex and insular cortex. Only the t-values of the positive significant cluster are shown. The black contour indicates the regions with the top 1% t-values across the whole brain. The t-values were interpolated and projected onto MNI coordinates for visualization purposes. The white contour indicates distractor-evoked neural activity, quantified as the top 1% inter-trial phase coherence (ITPC) in the post-distractor time window (i.e., 0 – 0.3 s) at 3 – 7 Hz (shown also in Fig. 2B). Bottom right panel shows centred perceptual sensitivity sorted by phase bins in the positive cluster at 3 Hz averaged across participants. Grey thin lines show individual centred perceptual sensitivity.

## Results

In the current electroencephalography (EEG) and behavioural study, we aimed at (1) uncovering the temporal fluctuations in distraction, and (2) exploring the relationship between such fluctuations and momentary neural phase at similar frequencies. To this end, we varied the onset time of an auditory distractor that was presented in-between two to-be-compared tones in a variant of a pitch discrimination task.

### Distractors interfere with pitch discrimination performance

To examine the potency of the distractors to distract, we compared participants’ sensitivity and criterion (response bias) of the pitch discrimination task between distractor-present and -absent trials. Participants were less sensitive to the pitch difference (*t_29_* = −8.11, *p* = <.001, Cohen’s d = −1.48), and had a more conservative response criterion (i.e., more “same pitch” responses; *t_29_* = 2.83, *p* = .008, Cohen’s d = 0.52) on distractor-present trials (Fig. 1B).

As the pitch difference presented for each participant differed depending on participant’s threshold in the adaptive task, we tested whether participants’ thresholds explain the susceptibility to distraction, which is quantified as the difference in behavioural sensitivity between distractor-absent and distractor-present conditions. Participants’ thresholds did not explain the degree of distraction (*t_28_* = −0.31, *p* = .76).

### Behavioural and neural measures of distraction co-fluctuate across time

Does the impact of distraction on neural activity and goal-directed behaviour exhibit fluctuations across time? To test this, we varied distractor onset time and examined whether behavioural (i.e., sensitivity; Fig. 2A, yellow) and neural measures of distraction (i.e., distractor-evoked ERP; Fig. 2A, blue) would show modulations at frequencies up to 8 Hz. To examine temporal fluctuations of distraction, we used linear mixed-effects models with sine- and cosine-transformed distractor onset time as predictors to model the behavioural and neural time courses as the outcome measures.

Fig. 2E and F show the spectral magnitude (0.5 – 8 Hz, 0.5-Hz steps) resulting from linear mixed models on detrended perceptual sensitivity (Fig. 2C, yellow) and detrended ERP amplitude (Fig. 2C, blue), respectively. Statistical significance was derived by testing empirical spectral magnitude against the 95^th^ percentile of a permutation distribution, which was derived from shuffling the behavioural and neural time courses, respectively, 5,000 times (see Methods for details).

At the behavioural level, distractor onset time modulated sensitivity below 5 Hz. At the neural level, distractor onset time modulated the distractor-evoked ERP at 4 and 5 Hz. However, only the 5 Hz spectral peak in sensitivity (Fig. 2E) stayed significant after FDR correction for all the frequencies (fluctuations below 2 Hz reflect slow trends in the data and are thus not discussed here).

Similar results were obtained in a control analysis, where temporal fluctuations in the behavioural and neural time courses in distractor-present trials were compared against distractor-absent trials (instead of permuted distractor-present trials; Fig. S6).

If these periodic neural dynamics serve as the basis for the apparent behavioural fluctuations, we should observe the synchronization of the behavioural and neural time courses by a common rhythm. To test this, we also examined the co-modulation of sensitivity and distractor-evoked ERP by distractor onset time. We first calculated the cross-correlation coefficients of the behavioural and neural time courses for individual participants (Fig. 2D). We then ran a linear mixed model with the cross-correlation coefficient as the outcome measure and sine- and cosine-transformed time lag as predictors.

Fig. 2G shows that sensitivity and distractor-evoked ERP are co-modulated at 3.5 and 5 Hz. At lag 0, there was a negative correlation between sensitivity and the distractor-evoked ERP, consistent with the notion that stronger distractor encoding (i.e., larger distractor-evoked ERP) corresponds to worse task performance (i.e., lower sensitivity). T-tests against zero on the (Fisher-z transformed) correlation coefficients across participants show that this correlation at time lag 0 was close to statistical significance (Pearson’s *r*: *t_29_* = −1.85, *p* = 0.08, mean Pearson’s *r* = −0.08; Spearman’s *r*: *t_29_* = −2.13, *p* = 0.04, mean Spearman’s *r* = −0.10).

As a control analysis, the same analysis pipeline was run on the data in the distractor-absent condition by randomly assigning a “distractor onset” for each distractor-absent trial, which did not reveal any significant co-modulation (Fig. S7): Neither time courses of sensitivity nor distractor-evoked ERP were modulated by randomly assigned distractor onset time; time lags did not modulate the cross-correlation of these two at any frequency. The temporal co-fluctuations of behavioural and neural measures of distraction at 3 – 5 Hz in distractor-present trials may be a manifestation of an underlying distractibility rhythm, which we probed into next.

### Pre-distractor neural phase in inferior frontal/insular cortex explains distraction

If the human brain hosts an endogenous rhythm that underlies distractibility dynamics, the neural state prior to distractor onset should explain the participant’s momentary vulnerability to interference by a distractor. To test this, we studied how pre-distractor neural phase relates to our previously established proxy of distraction, that is, behavioural sensitivity. We asked when in time and in which brain network(s) such an endogenous rhythm underlying distractibility would show up.

We employed source-projected EEG time courses to extract the quadratic relationship between the binned pre-distractor neural phase and perceptual sensitivity. For each trial (Fig. 3A), we first transformed a source-projected EEG data segment (0.4 s; sliding window) into the frequency domain using FFT. We then extracted neural phase for a given frequency (Fig. 3B). To calculate sensitivity sorted by phase bin, we first sorted the trials according to their phase values into 9 phase bins of equal size, followed by calculation of perceptual sensitivity for each bin (see Fig. S8 for individual participants’ sensitivity by phase bin). The same procedure was repeated for a range of frequencies (2.5 – 8 Hz, 0.5-Hz steps) and time windows (−0.3 – 0.3 s around distractor onset, 0.05-s steps). A cluster-based permutation test with the dimensions time, frequency, and voxels, wherein the quadratic fit was tested against zero, revealed a positive significant cluster (Fig. 3C; the same analyses with 7, 8, and 10 phase bins yielded comparable clusters across all dimensions and comparable statistical significance). The quadratic modulation of sensitivity by neural phase at 2.5 – 7.5 Hz was most prominent in the left insular and the inferior frontal cortices in the time window spanning ∼300 ms before distractor onset (cluster p-value = .026, two-tailed; see Fig. S9 for brain surface plots from other viewing angles).

To test whether the significant cluster overlaps with sources of auditory-evoked activity in auditory cortex regions, we compared its source with the source of distractor-evoked intertrial phase coherence (ITPC) at 3 – 7 Hz (shown also in Fig. 2B, bottom panel). Importantly, although the two effects were localized in proximal cortical regions (Fig. 3C, bottom panel), their core regions were mostly non-overlapping.

For control, we conducted the same analysis on the distractor-absent trials, which revealed no significant cluster (Fig. S10). We also tested the relationship between the pre-distractor neural phase and the post-distractor neural measure of distraction (i.e., 25-Hz amplitude of the distractor-evoked ERP), which did not reveal a significant effect (Fig. S11). Lastly, we tested whether there is also a quadratic relationship between pre-tone 2 neural phase and behavioural sensitivity. While there was no significant cluster in the distractor-present condition (all *p* > .06), a significant positive cluster was found in the distractor-absent condition (*p* = .01), but in different neural regions (i.e., left lingual gyrus and right inferior frontal cortex) from the left insular/inferior frontal origins found for distractibility dynamics. Importantly, in both distractor-present and distractor-absent trials, there was no significant cluster in the left insular or inferior frontal cortex, suggesting that pre-tone 2 neural phase in these regions does not explain fluctuations in behavioural sensitivity (Fig. S12).

## Discussion

The current study aimed to unravel the temporal dynamics of distractibility, using a pitch discrimination task with auditory distractors. The eventual degree of distraction and the neural processing of distractors were respectively quantified by distractor-evoked performance detriments and neural responses in the human electroencephalogram (EEG). We made a series of interesting observations.

First, the ∼3 – 5 Hz fluctuations of behavioural sensitivity across distractor onset time urged for the question whether the same fluctuations are observed in the human brain’s response to distractors. Consistently, we found that the distractor-evoked neural response covaries with behavioural sensitivity at similar frequencies. Second, while behavioural sensitivity and the distractor-evoked neural response might partly reflect post-perceptual processes (such as distractor suppression), we asked whether the brain hosts an endogenous oscillation that shapes the momentary state of distractibility. Confirming this, we found that pre-distractor neural phase in left inferior frontal/insular cortex explained rhythmic fluctuations in the momentary degree of distraction.

These major findings support the notion that temporal fluctuations in distractibility on a subsecond time scale can be explained by slow neural oscillatory dynamics in a cortical network beyond the auditory cortex.

### The proneness to distraction is inherently dynamic

The current study sheds light on the dynamics of distractibility, which is an important factor often neglected in previous attention research on distraction and suppression. The ultimate degree of detriment that a distractor will cause depends on two endogenous factors: the momentary proneness to distraction (i.e., distractibility) and the ability to suppress a distractor (i.e., distractor suppression). On the one hand, research on distractor suppression often did not disentangle the active suppression of distractors (Schneider et al., 2021) from variations in distractibility. On the other hand, research on distractibility rather treated it as an individual characteristic that, if at all, only changes on a slow temporal scale such as within an experimental session (Forster & Lavie, 2014) or across developmental stages (Kannass et al., 2006). The temporal trajectory of distractibility on a faster, subsecond, time scale had hitherto been left unknown.

With distractor-evoked behavioural and neural measures, we were able to encapsulate the temporal trajectory of distraction, which fluctuates on a subsecond temporal scale consistent with the rate of rhythmic sampling in attention (Fiebelkorn et al., 2013; Ho et al., 2017; Kubetschek & Kayser, 2021; Landau & Fries, 2012) and working memory (Cruzat et al., 2021; Schmid et al., 2022; ter Wal et al., 2021). With analysis of pre-distractor neural oscillatory phase, we were able to trace this distractibility back to a slow neural oscillatory fluctuation in inferior frontal and insular cortex (see below for an in-depth discussion). Participants could not anticipate whether or when the distractor would occur, thereby not being able to engage in preparatory suppression of the upcoming distractor (Geng, 2014). The combined analysis of pre-distractor neural phase and of post-distractor neural and behavioural measures complementarily elucidates how the brain alternates between states of higher and lower distractibility. These insights are essential for the inclusion of an explicit account of distraction in models of attention in psychology and neuroscience.

Fluctuations of distractibility at 3 – 5 Hz in the current study unveil the dynamic nature of attention, which was underappreciated in the static spotlight metaphor of attention (Posner et al., 1980). The attentional sampling of to-be-attended external stimuli (Fiebelkorn et al., 2013; Ho et al., 2017; Kubetschek & Kayser, 2021) or internal memory representation (Cruzat et al., 2021; Schmid et al., 2022; ter Wal et al., 2021) has been shown to exhibit temporal fluctuations at similar frequencies. The waxing and waning of attentional sampling may index inter-areal coordination between the attentional network and the sensory areas of the brain (Dugué & VanRullen, 2017), which is associated with the alternation between stronger and weaker attentional sampling over time (Fiebelkorn & Kastner, 2019). With much evidence on the temporally dynamic nature of the attentional spotlight, however, there is a lack of theoretical foundation for the inherent dynamics of cognition outside of this spotlight (Lui & Wöstmann, 2022). With the observed fluctuations of distractibility in the theta frequency range, an extension of the existing theory of dynamic attentional sampling to temporally dynamic distraction is warranted.

While our results demonstrate that distractibility exhibits temporal fluctuations, they do not reveal whether such fluctuations are independent of the fluctuations found in the attentional sampling of memory content. Participants in the current study had to maintain the memory representation of the pitch of tone 1 during a trial. The theta fluctuations found in the current study thus may represent the sampling of the internal representation of tone 1, with higher distractibility hypothetically occurring during the phase of reduced sampling of the memory representation. Alternatively, observed theta fluctuations may represent independent fluctuations in the proneness to distraction. Previous neuroimaging studies found that the suppression of distracting inputs may be independent of the sampling of attended inputs (Noonan et al., 2016; Schneider et al., 2018; Wöstmann et al., 2019). Future investigations may manipulate both the target and distractor onset time to examine the relationship between the temporal fluctuations underlying attentional sampling and distractibility.

Of note, as the main analysis approach used here (comparing empirical time courses to time courses that were shuffled in time) does not distinguish between periodic and aperiodic temporal structure (Brookshire, 2022), we are careful to conclude from the respective results alone that distractibility is rhythmic. However, it does not negate the possibility that there is a periodic temporal structure in distractibility. The premise of rhythmic cognition is that the apparent fluctuations of performance reflect the periodic orchestration between brain regions (Fiebelkorn & Kastner, 2019). In addition to fluctuations in behavioural performance, neural evidence is therefore essential to elucidate the rhythmicity of cognition (Fiebelkorn, 2022; Wöstmann, 2022). The current study shows a correspondence between slow neural oscillatory phase and behaviour (using an analysis approach that does not employ shuffling-in-time), consistent with the notion that distractibility is rhythmic. Future advancements in the analysis approach to directly test the periodicity in cognition will further strengthen our understanding of the distractibility dynamics.

While not all spectral peaks in the analyses survive FDR correction, all analyses with different outcome measures (i.e., behavioural sensitivity, distractor-evoked ERP, and cross-correlation), different contrasts (i.e., compared against permutation distribution or against distractor-absent condition), and with different analysis methods (i.e., linear mixed model with sine- and cosine-transformed distractor onset time or fast-Fourier transform) show consistent peaks between 3 and 5 Hz.

### Neural dynamics of distractibility originate in inferior frontal/insular cortices

The localisation of neural phase effects underlying distractibility dynamics beyond auditory cortex regions might suggest that the proneness to distraction is supra-modal. In research on visual distraction, brain regions in frontal and parietal cortices have been associated with distractor interference in lesions (Chao & Knight, 1995) or transcranial magnetic stimulation (Kanai et al., 2011; Wais et al., 2012) studies. The functional connectivity between the left inferior frontal cortex and hippocampus is associated with the disruptive influence of task-irrelevant visual distraction on working memory (Wais et al., 2010). While the current study examined distractibility in the auditory modality, the neural origins found here overlap with previous research on distraction in the visual modality.

The observed relationship between perceptual sensitivity and the inferior frontal/insular theta phase suggests that fluctuations in distractibility may be related to the cognitive control of working memory. The left inferior frontal cortex is assumed to be critical to the resolution of competition between the maintenance of goal-relevant information and the interference from external distraction (Irlbacher et al., 2014; Tops & Boksem, 2011; Wais et al., 2012). The anterior insula is theorised as a gatekeeper to the brain regions responsible for goal-related cognitive control (Molnar-Szakacs & Uddin, 2022), and is part of the ventral attention system (Eckert et al., 2009). Specifically, the insular cortex may support the switching between networks important to internally directed and externally directed cognition, respectively (Uddin, 2015). The frontal theta rhythm is associated with cognitive control (Berger et al., 2019; Cavanagh & Frank, 2014; Kamarajan et al., 2004) and the prioritization of relevant memory representation (Riddle et al., 2020). Taken together, theta oscillations in the inferior frontal and insular cortices may reflect the orchestration of the cognitive control system to maintain the internal memory representation and suppress potentially distracting external inputs.

We did not observe a corresponding phasic relationship between pre-tone 2 neural phase and behavioural sensitivity in the current study. Nevertheless, the absence of evidence here should not be taken as the evidence of absence as we did not manipulate target onset time after phase reset, which is crucial for unveiling any dynamics in auditory attentional sampling (Ho et al., 2017; Zoefel & Heil, 2013). To further understand the relationship between distractibility dynamics and the dynamics in attentional sampling, future studies should manipulate the onset time of targets and distractors to examine if attentional sampling and distractibility fluctuate at similar frequencies with different phases.

Against what might have been expected, pre-distractor theta phase did not predict fluctuations in the distractor-evoked neural response (Fig. S11). There are two salient reasons why the present data do not bear out a direct correspondence between pre-distractor neural phase and post-distractor neural response. First, the distractor-evoked ERP may not only reflect endogenous distractibility, but also other cognitive operations that contribute to the final degree of distraction, such as reactive suppression (Feldmann-Wüstefeld & Vogel, 2019; Hickey et al., 2009; B. Wang et al., 2019) or stimulus prediction (Volosin & Horváth, 2014). Distractibility dynamics may thus only account for a small amount of variance in the distractor-evoked ERP at 25 Hz. Second, in contrast to the distractor-evoked ERP, the neural origins of the distractibility dynamics rest in regions beyond the auditory cortex. The involvement of non-auditory cortical regions suggests that fluctuations in distractibility may arise from higher-order cognitive control processes, which affect behavioural sensitivity but are not directly related to distractor-evoked auditory responses.

## Conclusions

The present study demonstrates that human proneness to distraction is not uniformly distributed across time but fluctuates on a subsecond timescale in cycles of ∼3 to 5 Hz. In the brain, time windows of higher distractibility are coined by stronger neural responses to distractors. Furthermore, slow neural phase in left inferior frontal/insular cortex regions explains fluctuations in distractibility. These results unravel the temporal dynamics of distractibility and thereby help explain human processing of an abundant kind of stimulus in increasingly complex environments, that is, irrelevant and distracting input.

## Data availability

Raw data are available from the corresponding authors upon request. Pre-processed behavioural and neural responses as a function of distractor onset time will be made available online upon publication.

## Author contributions

Conceptualization, T.K.L., J.O., and M.W.; methodology, T.K.L., J.O., and M.W.; investigation, T.K.L.; formal analysis, T.K.L., J.O., and M.W.; writing – original draft, T.K.L. and M.W.; writing – review & editing T.K.L., J.O., and M.W.; funding acquisition, M.W.

## Acknowledgments

We thank Frauke Kraus and Malte Naujokat for their help in data collection.

---

**Fig. S1.**
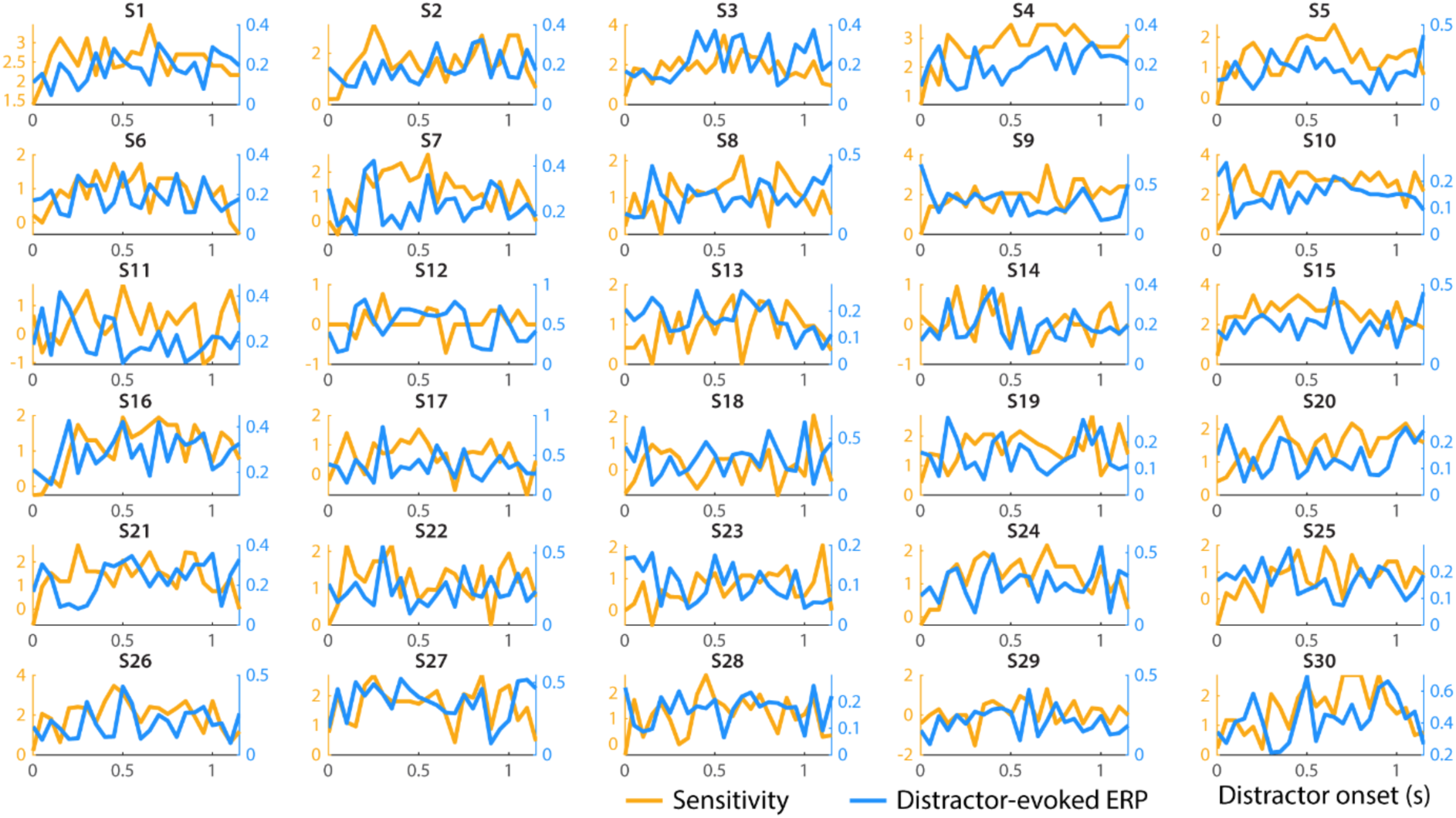
Individual time courses of raw sensitivity (yellow) and distractor-evoked ERP (25-Hz amplitude of distractor-evoked ERP; blue).

**Fig. S2.**
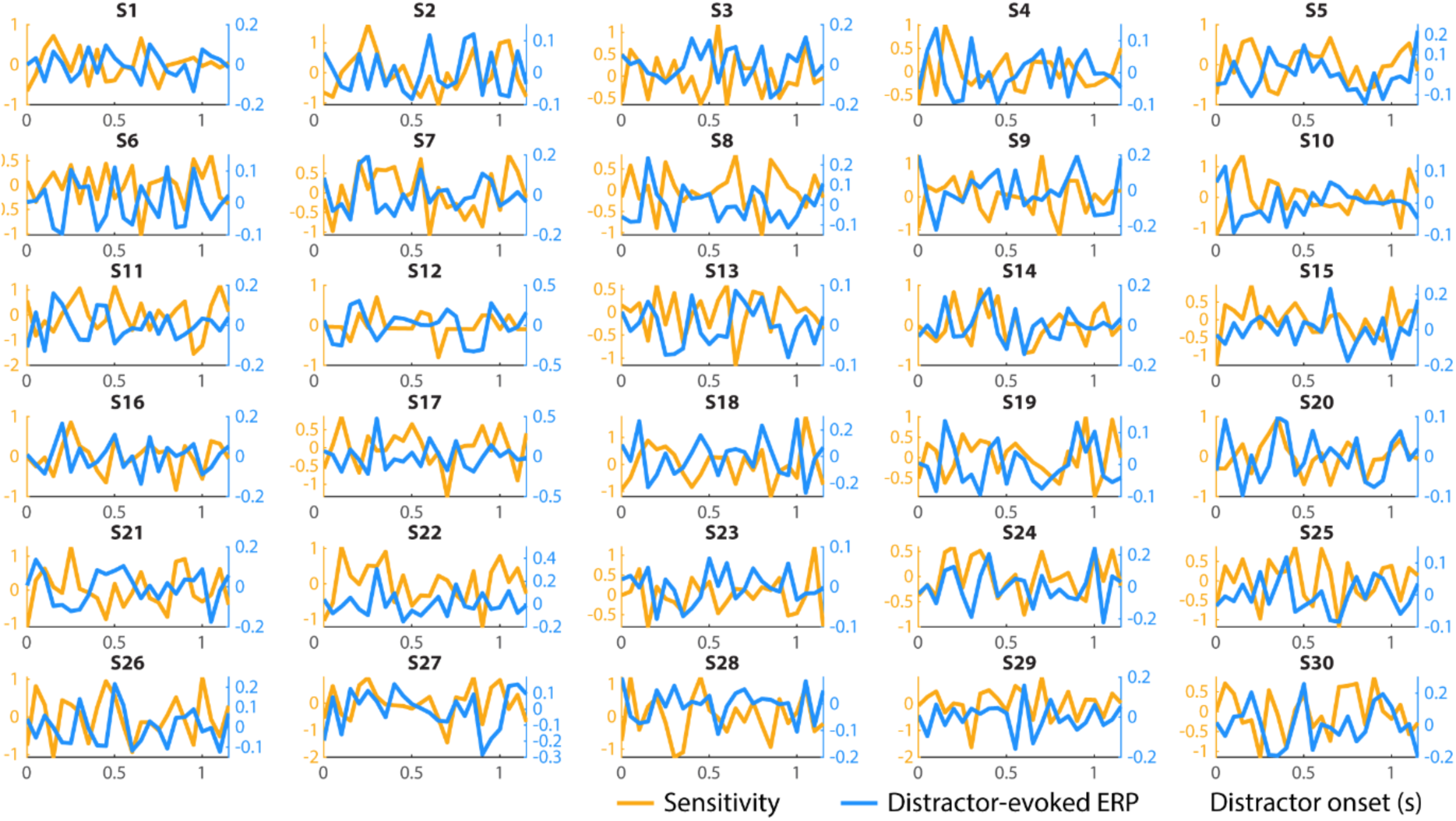
Detrended (quadratic trend removed) sensitivity (yellow) and distractor-evoked ERP (blue) individual time courses.

**Fig. S3.**
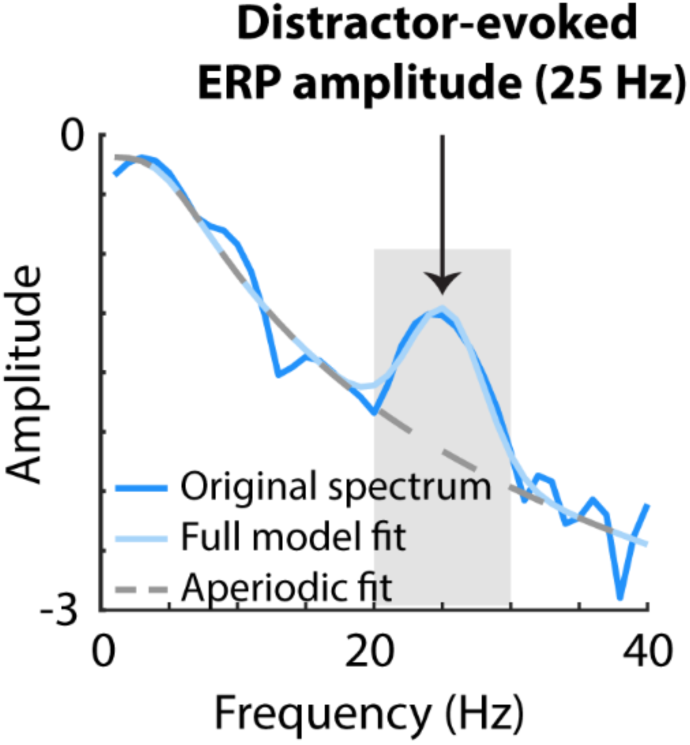
Frequency spectrum of the distractor-evoked ERP amplitude. Blue line shows the original frequency spectrum obtained by FFT. Light blue line shows the periodic signal (light blue) on the frequency spectrum while grey line shows the aperiodic activity (obtained from the fooof toolbox). Shaded grey area shows the frequency band (i.e., 20 – 30 Hz) at which the distractor-evoked ERP in Fig. 2 was filtered.

**Fig. S4.**
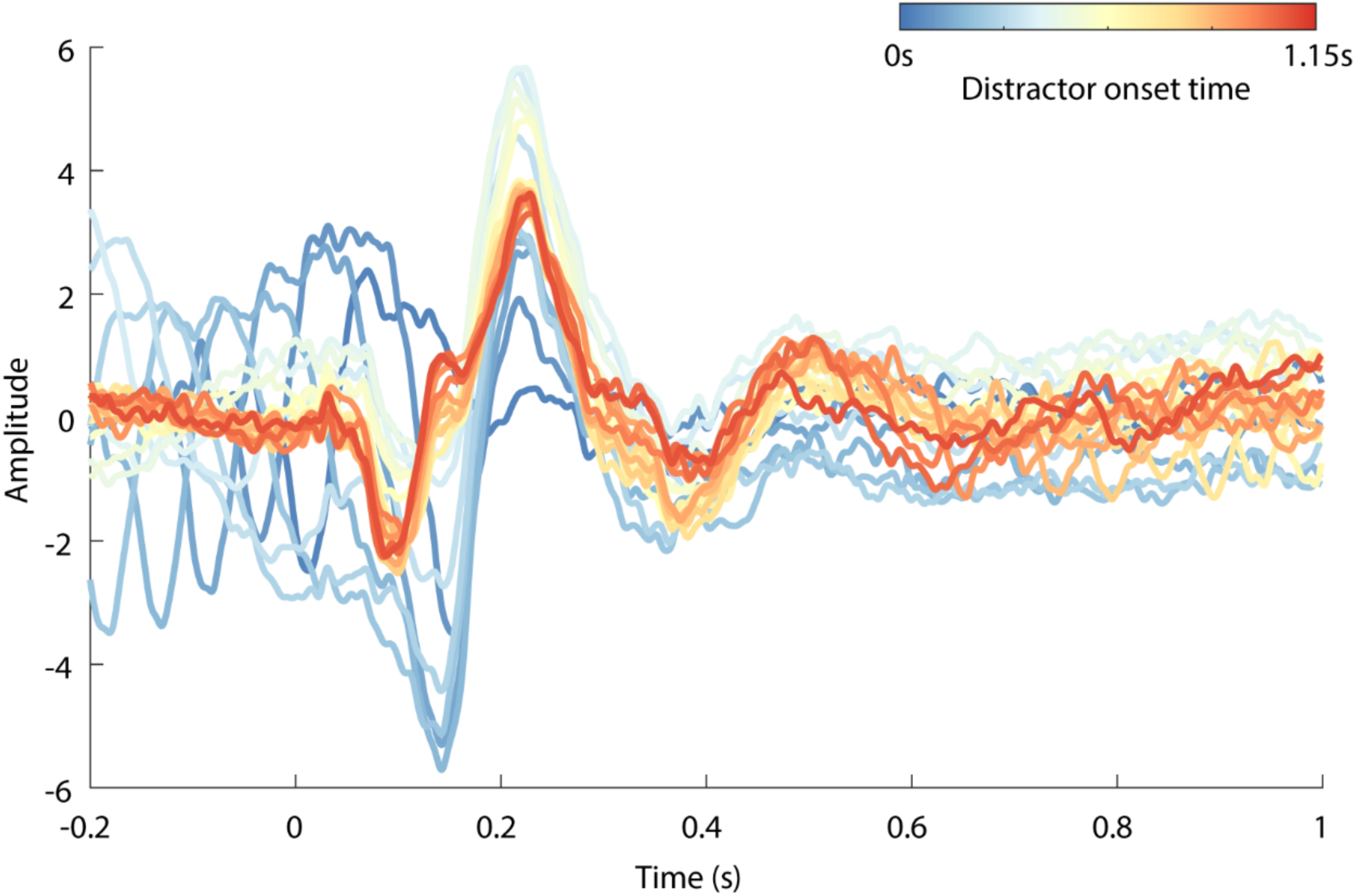
Distractor-evoked ERP waveforms, colour-coded for distractor onset time.

**Fig. S5.**
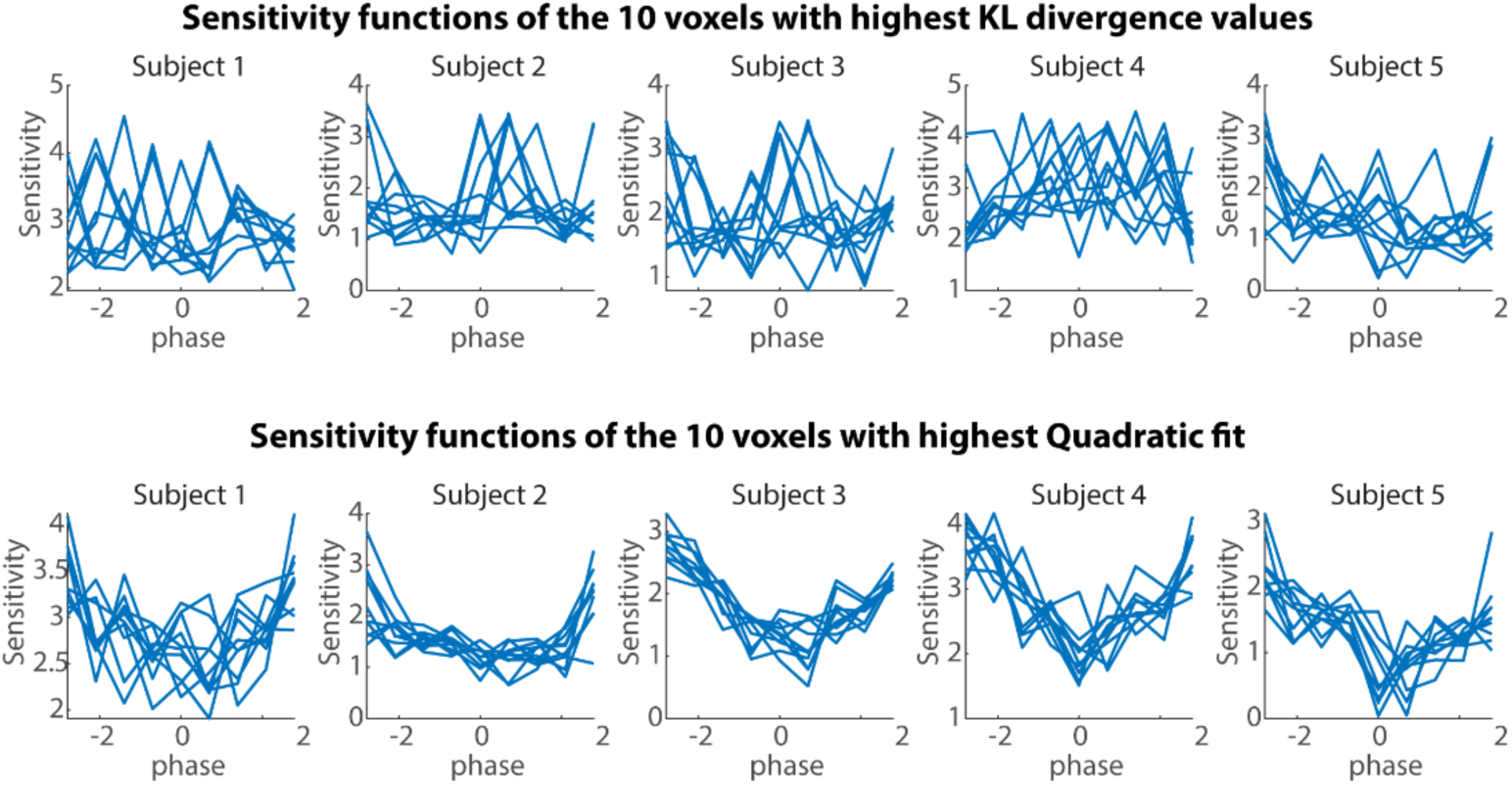
Sensitivity as a function of pre-distractor neural phase at the cluster peak effect (i.e., 3 Hz, −0.2 s relative to distractor onset) of 5 exemplary participants. Top row shows the 10 voxels with 10 highest KL divergence values. Bottom row shows 10 highest quadratic fit values. The sensitivity functions in the top and bottom panels belong to different voxels in the brain

**Fig. S6.**
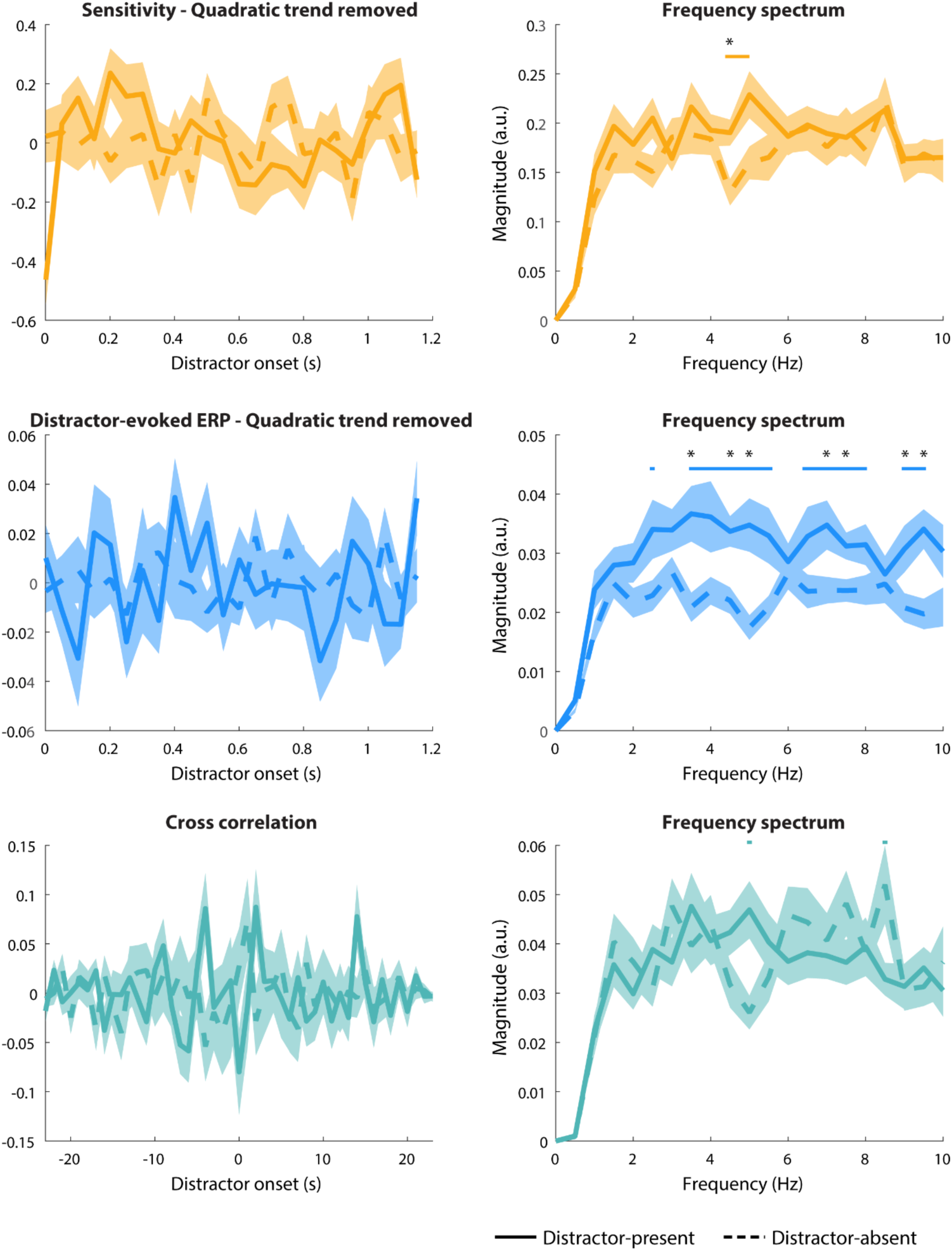
Comparison of the spectral magnitude between distractor-present and distractor-absent conditions. For behavioural sensitivity (top), distractor-evoked ERP (middle), and cross-correlation (bottom) time courses, we obtained the frequency spectra using FFT separately for distractor-present and -absent condition. For each frequency, we ran a dependent-samples t-test to compare the spectral magnitude between distractor-present and -absent conditions with correction of multiple comparisons (i.e., FDR correction). Left panels show the sensitivity (top), distractor-evoked ERP (middle), and cross-correlation (bottom) time courses for distractor-present (solid) and distractor-absent (dashed) conditions. Right panels show the averaged frequency spectra, derived from FFT on single-subject time courses. Shaded areas show ±1 SEM across individual participants. Horizontal lines show statistical significance when comparing the spectral magnitude of distractor-present versus-absent conditions (using dependent-samples t-tests) before FDR correction. Asterisks show statistical significance that remained significant after FDR correction. The statistical significance in sensitivity at 4.5 Hz remained significant after the correction of multiple comparison. The contrast between the distractor-present and -absent conditions on distractor-evoked ERP amplitude also show fluctuations in 3 – 5 Hz and 7 – 9.5 Hz after FDR correction.

**Fig. S7.**
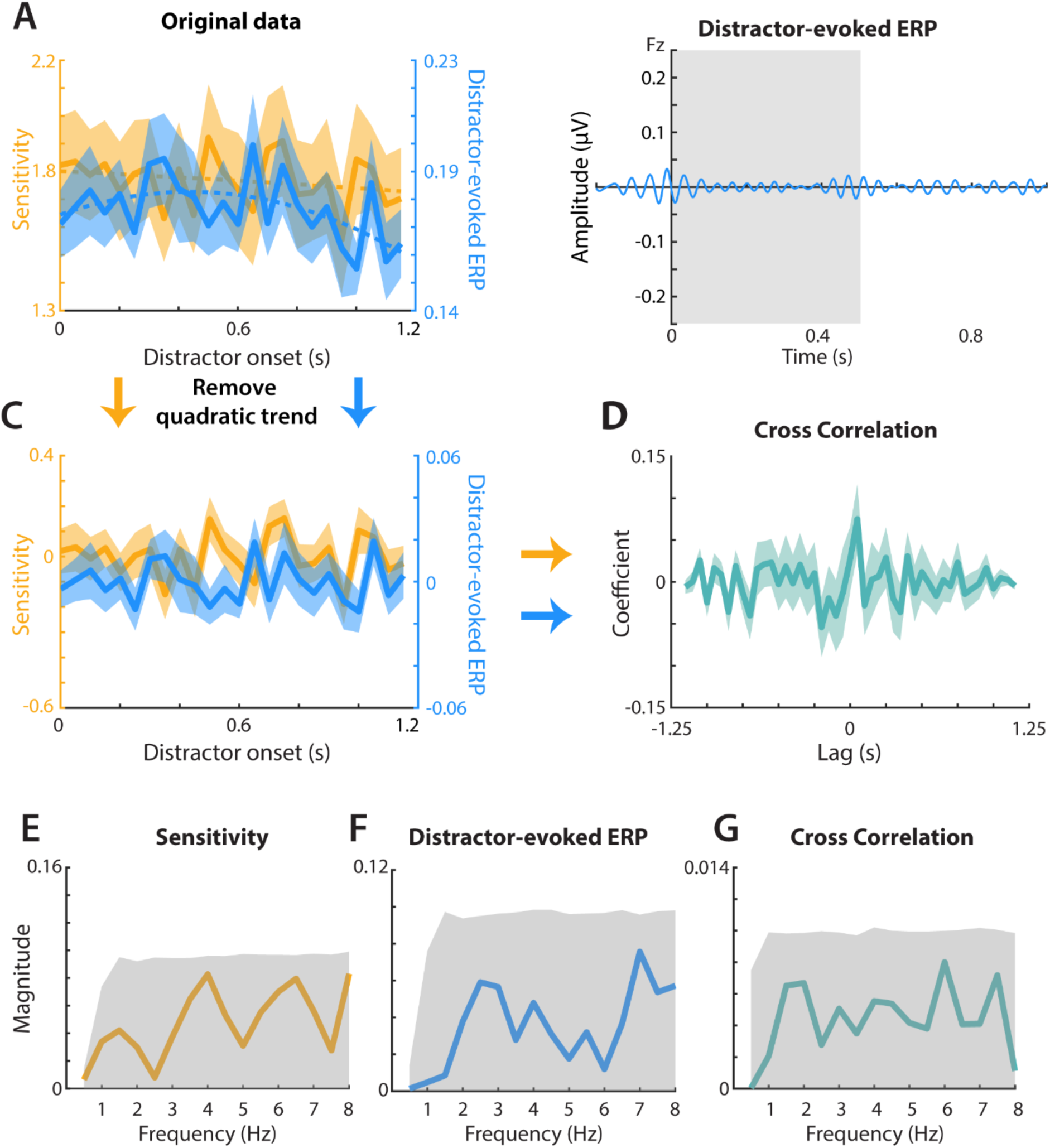
The same analysis pipeline as shown in Fig 2 applied to the distractor-absent condition.

**Fig. S8.**
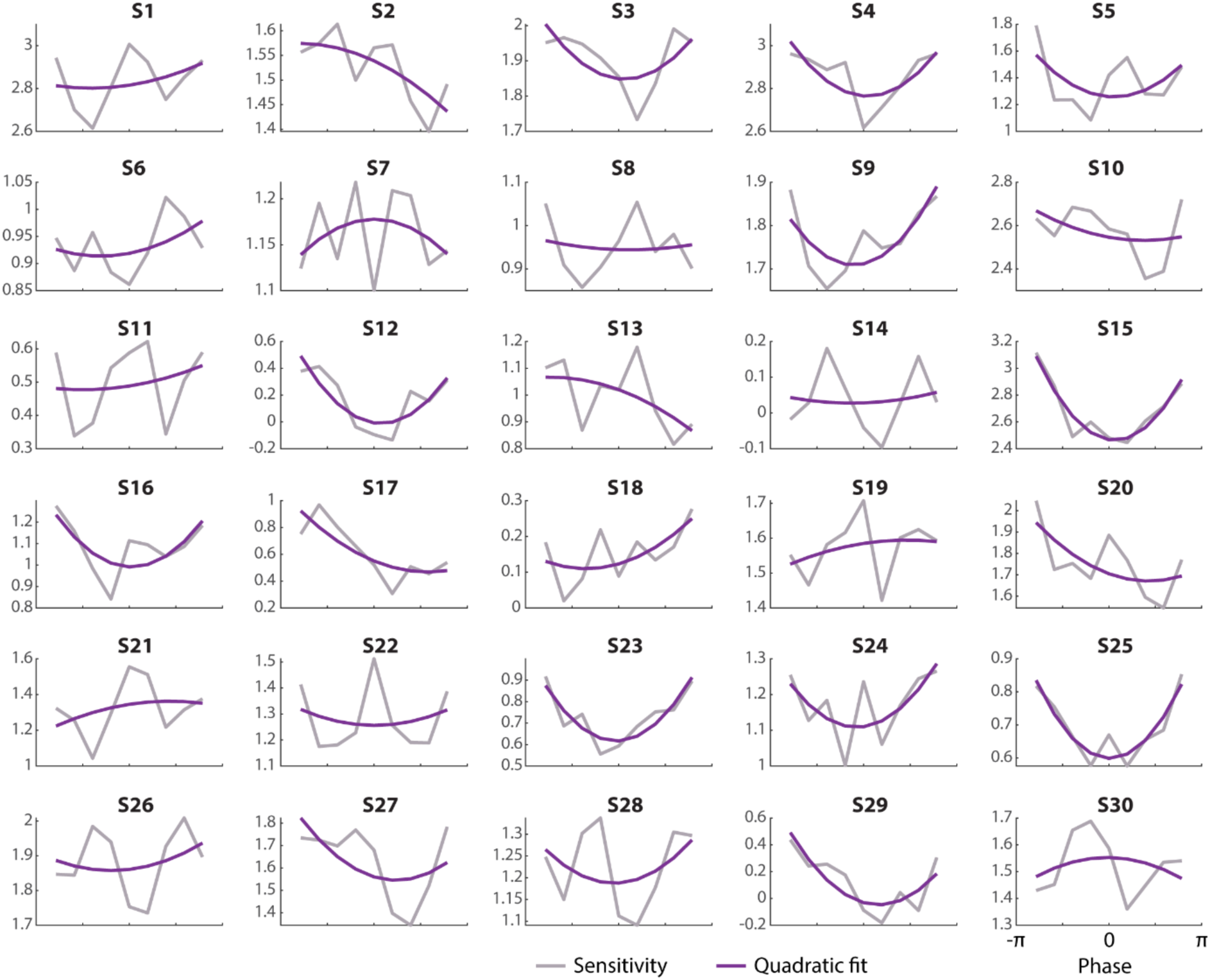
Individual sensitivity (grey line) and quadratic fit (purple line) across phase bins of the significant positive cluster at 3 Hz.

**Fig. S9.**
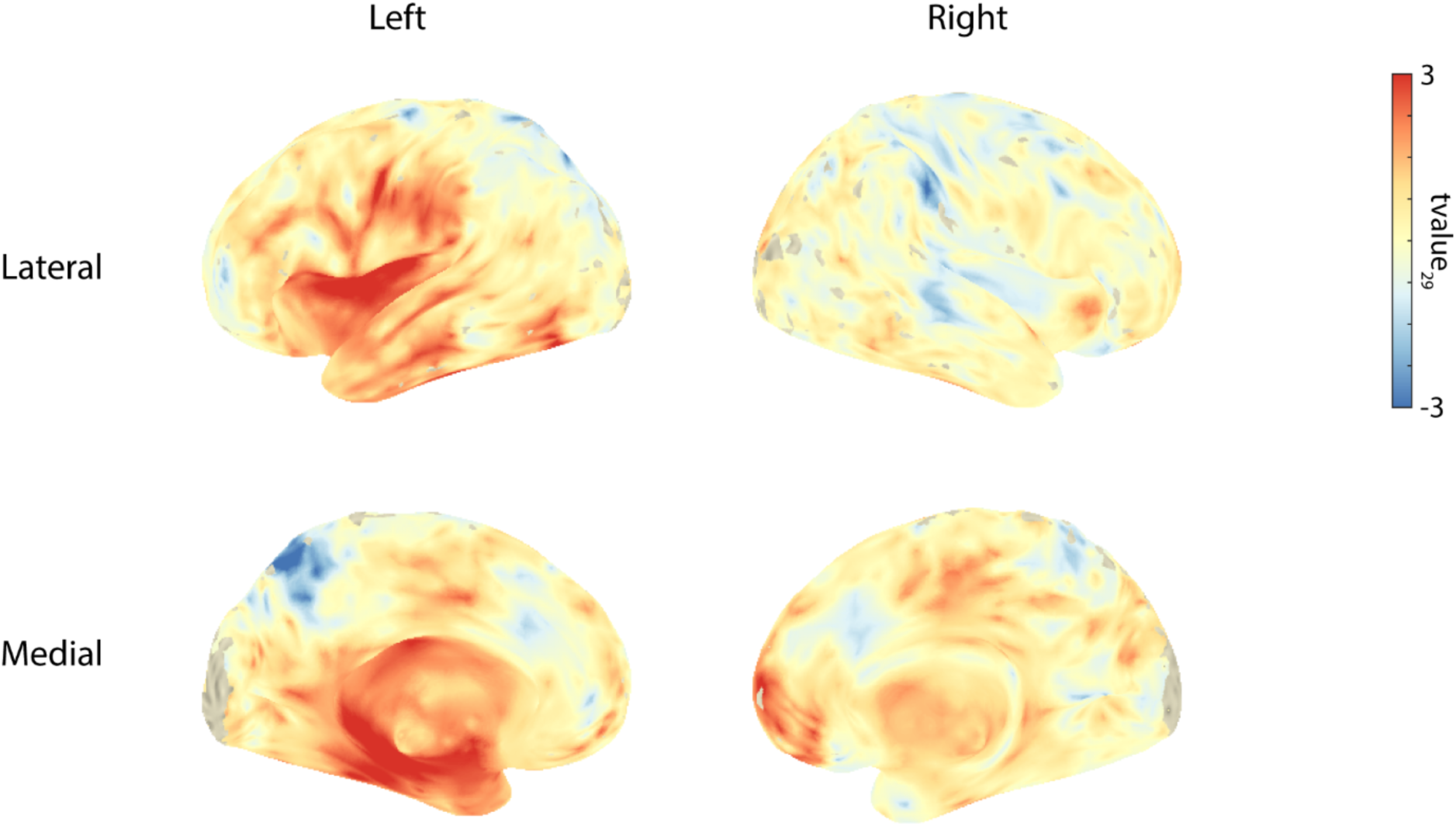
Brain surface plots of the cluster peak effect. Brain surface plots of the cluster peak effect (3 Hz; −0.2 s) from left lateral (top left), right lateral (top right), left medial (bottom left), and right medial (bottom right) views show t-values for the comparison of the quadratic fit of the sensitivity sorted by phase bins against zero. The t-values were interpolated and projected onto MNI coordinates for visualization purposes.

**Fig. S10.**
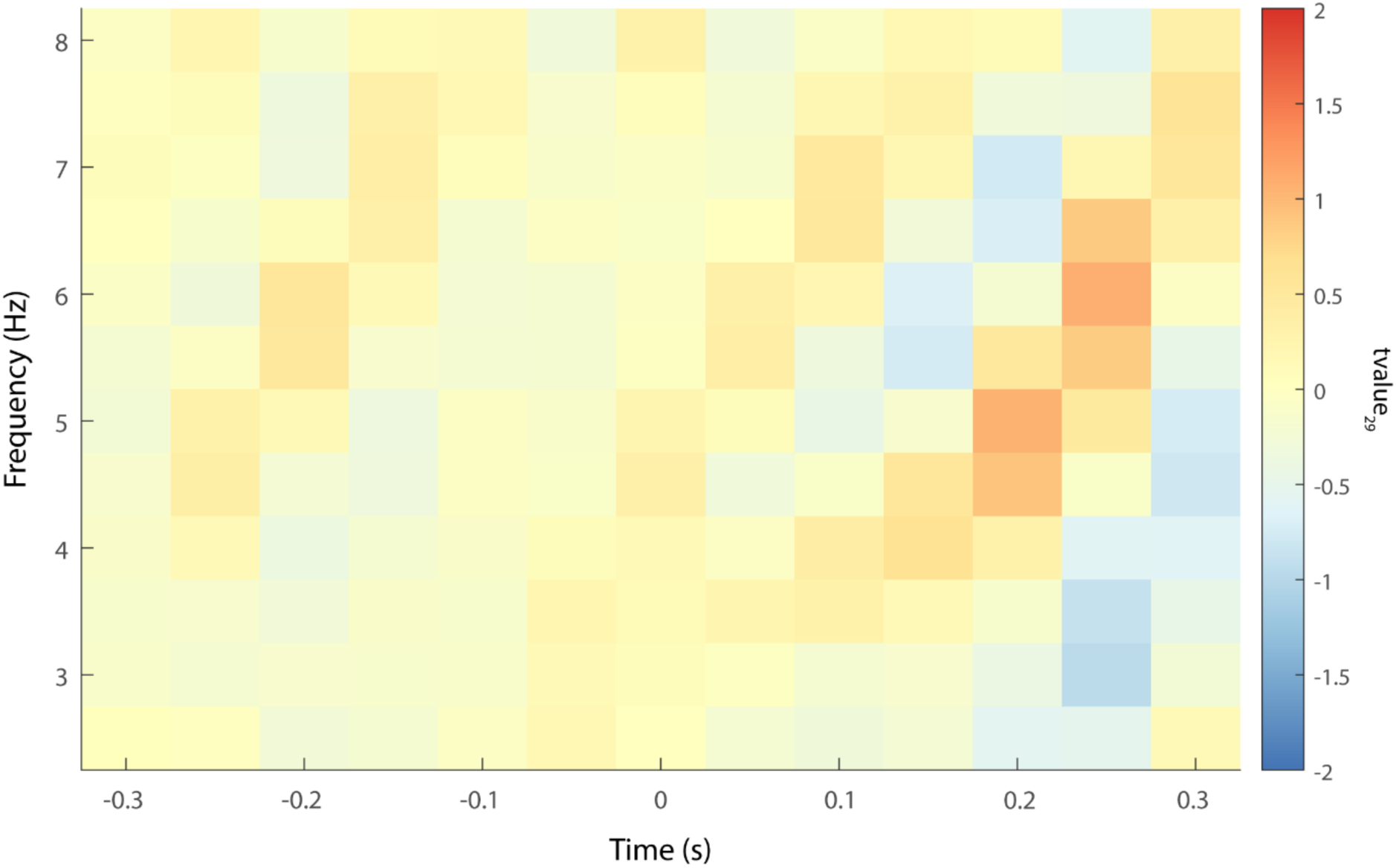
Cluster-based permutation test in the distractor-absent condition. Results of the cluster-based permutation test, across time windows and frequencies, on the quadratic relationship between neural phase and sensitivity for the distractor-absent condition. Figure shows t-values (df = 29) averaged across all the voxels belonging to the significant positive cluster in the distractor-present condition shown in Figure 3. No significant cluster was found in the distractor-absent condition.

**Fig. S11.**
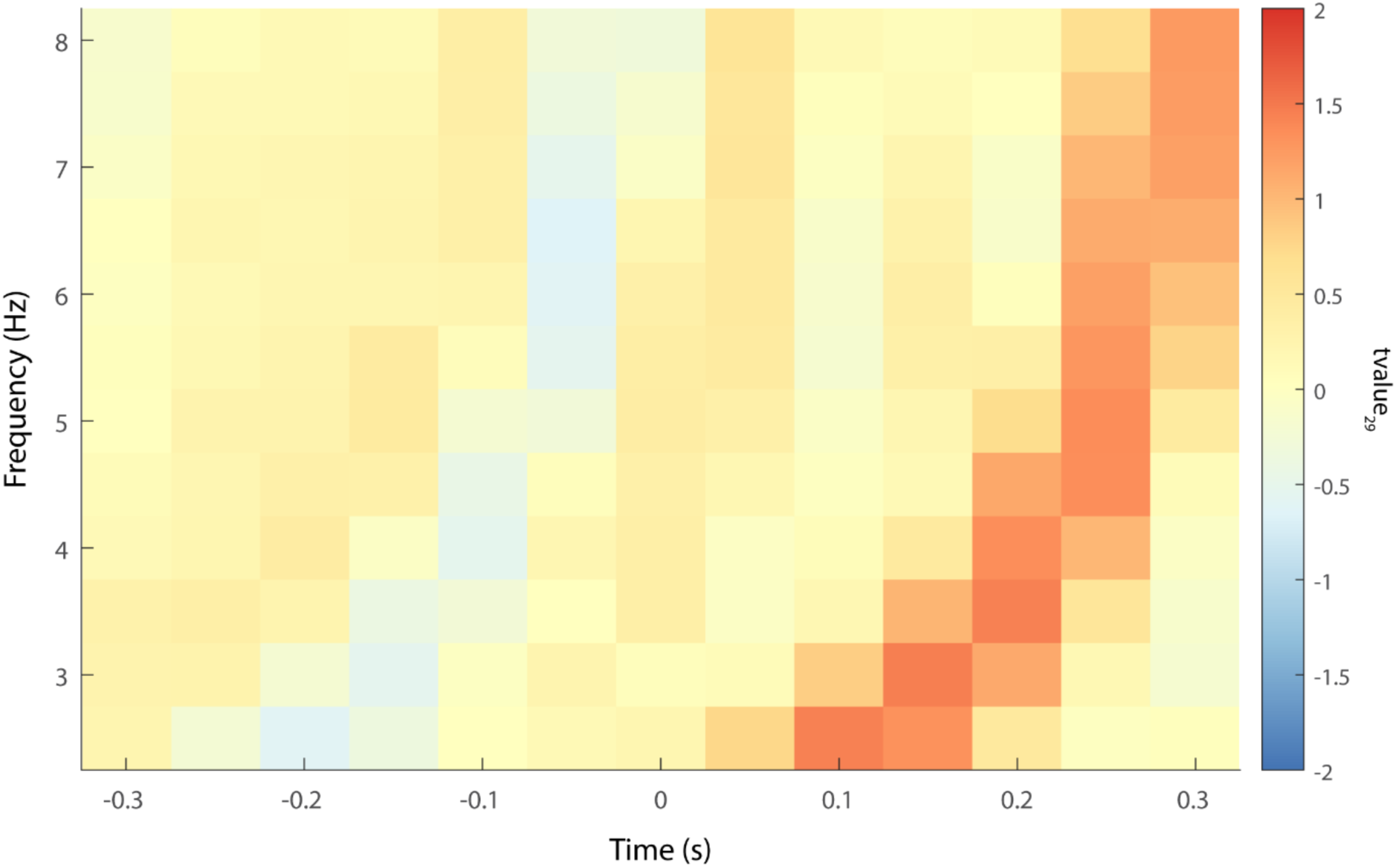
Cluster-based permutation test on distractor-evoked ERP amplitude. Results of the cluster-based permutation test, across time windows and frequencies, on the quadratic relationship between neural phase and distractor-evoked ERP amplitude at 25 Hz. Figure shows t-values (df = 29) averaged across all the voxels belonging to the significant positive cluster testing the quadratic relationship between neural phase and behavioural sensitivity shown in Figure 3. As expected, and not of main interest in the current study, a significant cluster in the post-stimulus time window (i.e., >0s) was found. More importantly, no significant cluster was found in the pre-distractor time window (i.e., <0 s).

**Fig. S12.**
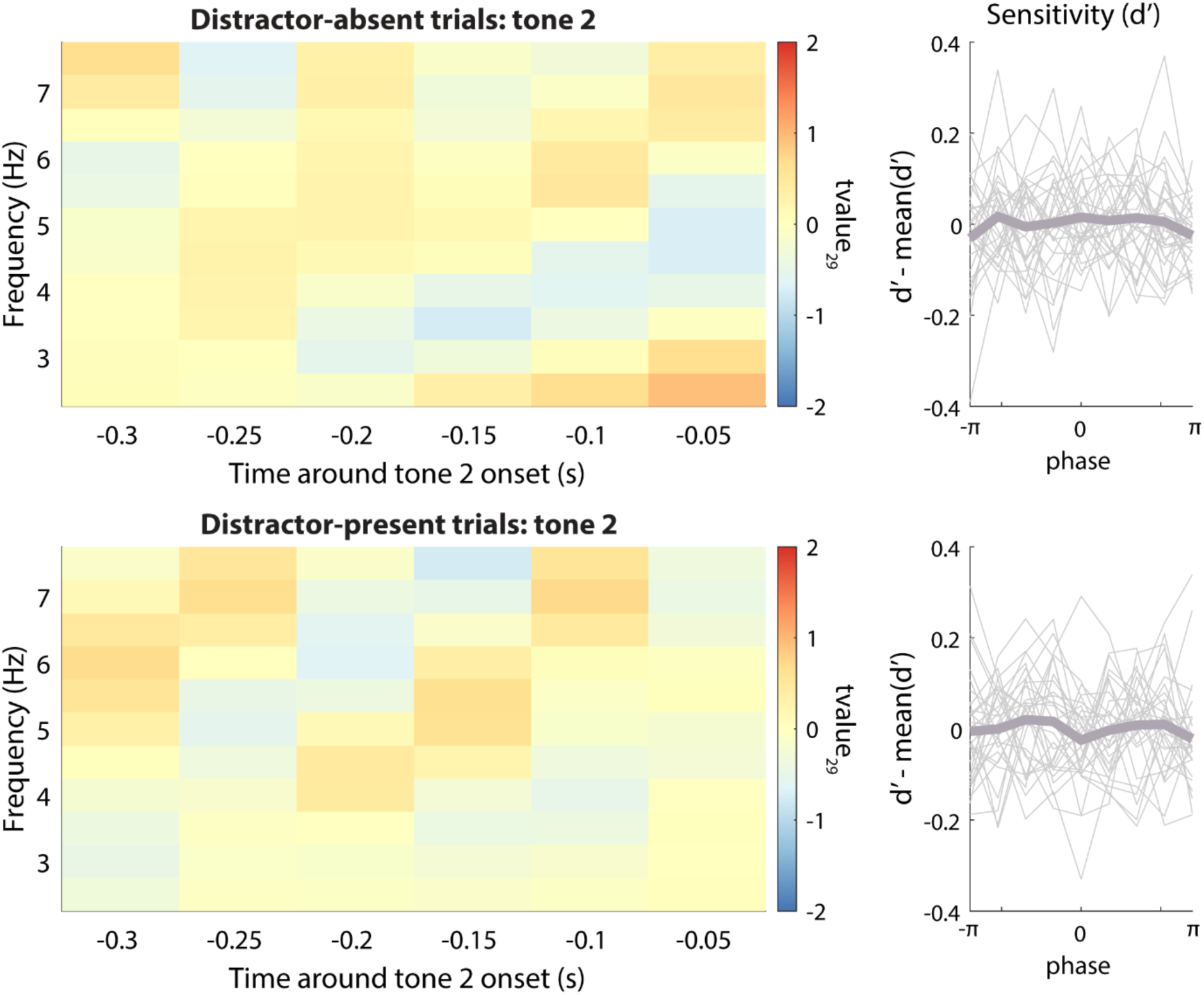
Control analysis of a possible relationship between pre-tone 2 neural phase and behavioural sensitivity. Results of the cluster-based permutation test, across time windows and frequencies, on the quadratic relationship between pre-tone 2 neural phase and sensitivity for distractor-absent (top panel) and distractor-present conditions (bottom), respectively. Left column shows t-values (df = 29) averaged across all the voxels belonging to the significant positive cluster testing the quadratic relationship between pre-distractor neural phase and behavioural sensitivity shown in Fig. 3 in the main manuscript. Right column shows the centred perceptual sensitivity sorted by phase bins in the same cluster at 3 Hz and −0.2 s averaged across participants. Grey thin lines show individual centred perceptual sensitivity.

